# Transcription profiling and functional analysis of spRNAs and their corresponding asRNAs in *Methanosarcina mazei*

**DOI:** 10.1101/2025.10.28.685009

**Authors:** B. Jordan, R. Gohil, D. Prasse, L. Hellwig, U. Repnik, S. Jiang, J. Yuan, R. A. Schmitz

## Abstract

Small proteins (sPs) encoded by small RNAs (sRNAs), the so-called small protein RNAs (spRNAs), and their corresponding antisense RNAs (asRNAs) have emerged as important regulatory elements in microbial stress adaptation, yet remain poorly characterized in archaea. In this study, we identified and characterized five sRNA/asRNA pairs in the methanoarchaeon *Methanosarcina mazei*. Expression analysis revealed condition-specific regulation of two of those pairs, spRNA23/asRNA93 and spRNA24/asRNA94, under oxidative, temperature, and osmotic stress. Overexpression of the corresponding small proteins, sP23 and sP24, revealed distinct physiological effects: both enhanced growth under high-salt conditions, while sP24 in addition supported cell size maintenance during osmotic stress, as confirmed by transmission electron microscopy. Consistently, overexpression of asRNA94 let to impaired growth under salt stress, suggesting its negative regulatory role towards sP24. The results also imply that the antisense RNA might be crucial for controlling sP24 expression under non-stress conditions. Bioinformatic analysis predicted one transmembrane domain in sP24. Membrane-association of sP24 was experimentally validated *in vitro* using a cell-free expression system, demonstrating sP24’s integration into lipid bilayers. Conservation analysis showed that spRNA24 appears specific for *M. mazei* strains potentially reflecting niche adaptation, while spRNA23 is widespread within *Methanosarcina*. These findings demonstrate that archaeal sRNAs can encode functional small proteins, often encoded together with their respective regulatory asRNA, that contribute to environmental stress responses, overall offering new insights into the role of sRNA-derived small proteins in archaea.

## 1 Introduction

The discovery of small non-coding RNAs (sRNAs) has significantly expanded our understanding of gene regulation in both prokaryotes and eukaryotes. Advances in RNA sequencing technologies, coupled with experimental and computational tools, have led to the identification of numerous sRNAs across bacterial and archaeal species (Babski et al., 2016; Babski et al., 2011; Ghandour & Papenfort, 2023; Heyer et al., 2012; Jäger et al., 2009; Papenfort & Melamed, 2023; Papenfort & Vogel, 2010; Straub et al., 2009). These regulatory RNAs are generally classified into three categories: trans-encoded sRNAs, cis-encoded antisense RNAs (asRNAs), and guide RNAs involved in RNA modification (Straub et al., 2009; Wagner & Romby, 2015). While the molecular functions of bacterial sRNAs are well documented, knowledge of their roles in archaea remains limited (Bernick et al., 2012; Jäger et al., 2012; Jaschinski et al., 2014; Märtens et al., 2013; Prasse et al., 2017; Prasse & Schmitz, 2018).

Beyond classical regulatory sRNAs, a subset of sRNAs has been identified that encode small proteins and are therefore termed small protein RNAs (spRNAs). These spRNAs contain small open reading frames (sORFs) that encode proteins typically below 70 amino acids in length (Aoyama et al., 2022; Burton et al., 2024; Hemm et al., 2020; Hobbs et al., 2011; Patraquim et al., 2020; Ramamurthi & Storz, 2014; Storz et al., 2014; Weidenbach et al., 2022). Due to their small size, such ORFs are often overlooked by conventional genome annotation pipelines. However, with advances in RNA-seq, Ribo-seq, and bioinformatic methods, an increasing number of sORFs have been predicted in microbial genomes (Chihara et al., 2022; Hör et al., 2020; Vazquez-Laslop et al., 2022). Although some of these small proteins have defined functions (Steinberg & Koch, 2021; Storz et al., 2014; Weidenbach et al., 2022; Yadavalli & Yuan, 2022), the majority remain uncharacterized (Schlesinger & Elsässer, 2022). Intriguingly, some spRNAs act as dual-function molecules, serving both as regulatory RNAs and as templates for small protein synthesis (Gimpel & Brantl, 2017; Raina et al., 2018; Vanderpool et al., 2011; Venkat et al., 2021).

Methanoarchaea are strictly anaerobic microorganisms characterized by methane production as their primary catabolic process (Burton et al., 2024; Deppenmeier et al., 1996; Hook et al., 2010; Liu, 2010; Thauer et al., 2008). Among them, the genus *Methanosarcina* stands out for its metabolic versatility, which enables adaptation to diverse environments. *Methanosarcina mazei* Gö1, a mesophilic archaeon within the order *Methanosarcinales*, exemplifies this adaptability. It can utilize a range of substrates like H₂/CO₂, methanol, methylamines, and acetate for methanogenesis (Deppenmeier et al., 1990; Deppenmeier et al., 2002; Hippe et al., 1979). Consequently, *M. mazei* contributes substantially to methane emissions, a key greenhouse gas (Ferry, 1999; Thauer, 1998), and also possesses the capacity for nitrogen fixation (Ehlers et al., 2002). The establishment of a genetic system for *M. mazei* has further enabled functional studies *in vivo* (Ehlers et al., 2011; Ehlers et al., 2005; Thomsen et al., 2022). In *M. mazei*, sRNA-mediated post-transcriptional regulation has been studied extensively under varying nitrogen conditions. A genome-wide differential RNA-seq (dRNA-seq) analysis revealed 248 differentially expressed sRNAs in response to nitrogen limitation, indicating a central role for sRNAs in nitrogen and stress response pathways (Jäger et al., 2009). Several regulatory sRNAs have been studied in detail, for which experimentally confirmed physiological functions have been observed, e.g. sRNA_154_, central nitrogen regulatory RNA (Prasse et al., 2017), sRNA_41_, involved in regulation of acetyl-CoA synthase/ CO dehydrogenase (Buddeweg et al., 2018), and sRNA_162_, involved in regulation in response to the different carbon sources (Jäger et al., 2012).

In the present study, we identified several spRNAs in *M. mazei* that are associated with corresponding antisense RNAs. Respective transcript levels were monitored under stress conditions, revealing that the spRNA23/asRNA93 and spRNA24/asRNA94 pairs exhibited the most significant expression changes in response to stress. Based on these findings, spRNA23 and spRNA24 were selected for detailed investigation, focusing on the expression and functional characterization of their encoded small proteins and their roles during stress adaptation. This revealed a potential beneficial role for sP23 and sP24 during salt stress with an indication that sP24 may play a critical role in maintaining normal cell physiology under those stress conditions.

## 2 Materials and Methods

### 2.1 Strains and plasmids

Strains and plasmids used in this study are listed in Table S1 and S2. Plasmids were transformed into *Escherichia coli* DH5α (Hanahan, 1983) or DH5α λpir (Miller & Mekalanos, 1988) as previously described (Inoue et al., 1990) and into *M. mazei* (Ehlers et al., 2005) by liposome mediated transformation with modifications (Gehlert et al., 2023; Thomsen et al., 2022).

### 2.2 Plasmid construction

All generated and uses plasmids are summarized in Table S2. For construction of overproduction mutants of the small proteins, plasmids containing the sequence of the predicted protein N-terminally fused to a His-tag under the control of the constitutive promotor P*mcrB* and the terminator *tmcr* (Guss et al., 2008) were delivered inserted in plasmid pEX A-128 by Eurofins genomics, Germany. asRNAs under control of P*mcrB* were delivered inserted in plasmid pEX K-168 by Eurofins genomics, Germany. The sequence was isolated from the plasmid by *SacI* and *KpnI* restriction and ligated into the shuttle vector pWM321 (Metcalf et al., 1997) or pRS1595 (Thomsen & Schmitz, 2022) linearized with *SacI* and *KpnI*. The resulting plasmid was transformed in *M. mazei* by liposome-mediated transformation as described (Gehlert et al., 2023; Thomsen et al., 2022).

The open reading frame of sORF24 was amplified via PCR using primer pair ORF24_1_f/ ORF24_rev and pRS1340 as template. Primer ORF24_1_f provides an artificial Shine-Dalgarno sequence for efficient translation in *E. coli* and an *EcoRI* restriction site. Primer ORF24_rev binds downstream of the T7 terminator of pRS1340 and provides an *XbaI* restriction site. The amplified fragment was cloned into a modified pBAD expression plasmid (Unoson & Wagner, 2008) using *EcoRI* and *XbaI* (FastDigest, Thermo Fisher Scientific, US) and T4 DNA ligase (New England Biolabs, US). The resulting plasmid was denoted p-sORF24. Site-directed mutagenesis of p-sORF24 was performed by PCR with primers listed in Table S2. After removal of template DNA via *DpnI* digestion (Thermo Fisher Scientific, US), plasmids were transformed into *E. coli* K-12 wild type MG1655. All constructs were confirmed by sequencing (Microsynth SeqLab, Germany). All used primers are listed in Table S3.

### 2.3 Growth of *M. mazei*

*M. mazei* cultures were grown under anaerobic conditions with a N_2_/CO_2_ (80 %/ 20 %, vol:vol) gas phase as described by Ehlers et al. (Ehlers et al., 2002), 1 ml of preculture was supplemented to closed bottles containing 50 ml minimal medium as previously described. The cells were incubated at 37 °C without shaking. For temperature stress, the cultures were incubated at 30 °C or 42 °C.

Growth experiments of *M. mazei* mutants were realized under different conditions in 50 ml minimal media. Complemented medium in large amounts (e.g. 0.5 L) was divided under anaerobic conditions into 50 ml aliquots and each inoculated with the same preculture to a turbidity at 600 nm (T_600_) of 0.05. Cultures were incubated at 37°C or 42°C and T_600_ was monitored over time. Taken samples were diluted with media when T_600_ exceeded 0.5. Temperature shift was performed by changing incubation temperature from 37°C to 42°C. Growth on high salt media was realized with minimal media containing 500 mM NaCl. In this case, precultures were already grown in 500 mM NaCl containing media. The average growth and respective standard deviation were calculated from three biological replicates.

### 2.4 RNA isolation

*M. mazei* wildtype cultures were grown as described above. The cultures were harvested at a turbidity of TD_600_ of approximately 0.5-0.6 by centrifugation at 2,455 × g and 4 °C for 30 min followed by RNA isolation with Rotizol (Carl Roth, Germany), following the manufactures protocol.

### 2.5 Northern Blot

Northern Blot analysis was performed using the protocol previously described (Jäger et al., 2009). RNAs were detected with 5′-^32^P labeled ssDNA oligonucleotide probes (Table S4). Quantification of the northern blots was performed with AIDA software (Raytest, Germany). The quantified intensity (PLS/pixel) was normalized to the intensity of the respective 5S rRNA. The x-fold regulation of one replicate was determined by calculating the relation of the intensity from the stress condition to the intensity of exponential growth phase of classical growth conditions. Three biological replicates were used to calculate the average fold regulation and the dedicated standard deviation.

### 2.6 Cell-free expression system with lipid sponge droplets

The cell-free expression with lipid sponge droplets was performed using the PURExpress from NEB as described previously (Cho et al., 2022; Bhattacharya et al., 2020; Jiang et al., 2024). Briefly, a thin lipid film was obtained by first mixing 42 μl of a single-chain galactolipid N-oleoyl β-D-galactopyranosylamine (GOA, 10 mM in methanol) with 100 μl chloroform, followed by evaporation to remove the solvent. Then, the lipid film was rehydrated with 21 μl solution including 2.66 μl of the non-ionic detergent octylphenoxypolyethoxyethanol (IGEPAL, 90 mM), 14 μl of the PURExpress Solution A, and 0.875 μl RNase inhibitor (NEB). Lipid sponge droplets were formed spontaneously in the rehydration process. The droplet solution (3 μl) was then combined with 1.5 μl PURExpress Solution B and 0.5 μl DNA template solution (150 nM) to assemble the reaction mix, an aliquot of which (3 μl) was then transferred into a lummox dish (Sarstedt) and sealed with a cover glass to prevent evaporation. Reactions were incubated at 37 °C on stage for three hours and observed using a Nikon Eclipse Ti-E inverted fluorescence microscope with a 40× objective.

### 2.7 Morphological analyses by electron microscopy

*M. mazei* cultures were cultivated for 72 h at 37°C in 50 ml media as described above. At TD_600_ of 0.4, cultures were fixed by adding 2% glutaraldehyde in 200 mM HEPES buffer to obtain the final concentration of 1% glutaraldehyde. The samples were incubated for 10 min at 37°C and for another 2 h at room temperature followed by centrifugation at 1811 x g for 10 min at 4°C. The pellet was resuspended in fresh fixative (1% glutaraldehyde in 200 mM Hepes pH 7.0) and incubated overnight at room temperature.

For morphological analysis, samples were imaged with a scanning electron microscope. 50 μl of a dense suspension of glutaraldehyde-fixed cells was transferred onto carbon and poly-L-Lysine coated cover slips. After 5 min, cover slips were washed with water, and samples incubated with 1% osmium tetroxide for 15 min, followed by 2% aqueous uranyl acetate for 15 min, and then dehydrated with an ethanol series 70-80-90-96-100-100%, each step for 10 min, followed by critical point drying (CPD 030, BAL-TEC AG). Dried cover slips were mounted on adhesive carbon tape and sputter coated to obtain a 5-nm platinum layer. Samples were imaged with a Sigma 300 VP (Zeiss) scanning electron microscope using a secondary electron (SE) detector and 2 kV accelerating voltage.

For whole-mount imaging with a transmission electron microscope, drops of cell suspensions were incubated on poly-L-lysine coated TEM grids for 10 min to let cells stick to the formvar support film, followed by washing with five drops of dH_2_O, and staining with1% uranyl acetate for 2-3 min. Stained grids were washed with three drops of H_2_O, blotted with filter paper and air dried before they were imaged with a CM10 transmission electron microscope (Philips), operated at 80 kV, using a MegaView III camera, and iTEM software (both Olympus Soft Imaging Solutions). Images were collected at 4,600x magnification with systematic uniform random sampling over a whole grid. For each sample group, three independent samples were analysed; for each sample 3 grids were imaged; on each grid at least 200 cells were imaged. Cell size was measured as the area of a cell profile using the Fiji software (Schindelin et al., 2012). Average values for each sample were used to test for statistically significant differences in the cell size between different genotypes within each of the two treatment groups (normal and 500 mM NaCl) using one-way ANOVA test. When the p-value corresponding to the F-statistic of one-way ANOVA was lower than 0.01, a post-hoc Tukey Honest Significant Difference (HSD) test was used to identify which of the three pairs of samples were significantly different from each other. Tests were run using an on-line test calculator (https://astatsa.com/OneWay_Anova_with_TukeyHSD/).

## 3 Results

### 3.1 Identification and Conservation of spRNAs and Corresponding asRNAs

A global transcriptomic analysis conducted by Jäger et al. (Jäger et al., 2009) using the 454 sequencing technique identified numerous sRNAs in *M. mazei*. Among these, transcripts containing putative sORFs were classified as spRNAs (Jäger et al., 2009). Several of those spRNAs, located in intergenic regions, have been found to be associated with asRNAs. Notably, five explicite spRNA/asRNA pairs were identified: spRNA09/asRNA60, spRNA10/asRNA61, spRNA23/asRNA93, spRNA24/asRNA94, and spRNA33/asRNA28.

The transcription profiles of these pairs under standard growth conditions were visualized using RNA-seq data (Illumina Miseq data) previously published by Nickel et al. (Nickel et al., 2013), depicting the asRNA reads in grey and spRNA reads in green (see Fig. 1). Predicted sORFs are represented as black-lined boxes; for spRNA33, three such boxes indicate multiple potential sORFs. The sense-to-antisense RNA transcript ratios vary across the pairs: spRNA09/asRNA60 and spRNA33/asRNA28 exhibit higher levels of sense transcripts, while spRNA10/asRNA61 and spRNA24/asRNA94 show approximately equal levels of sense and antisense transcripts. In contrast, the transcript ratio for spRNA23/asRNA93 is strongly shifted toward the antisense RNA, suggesting a potential regulatory mechanism that enhances the degradation of spRNA23.

**Figure 1:**
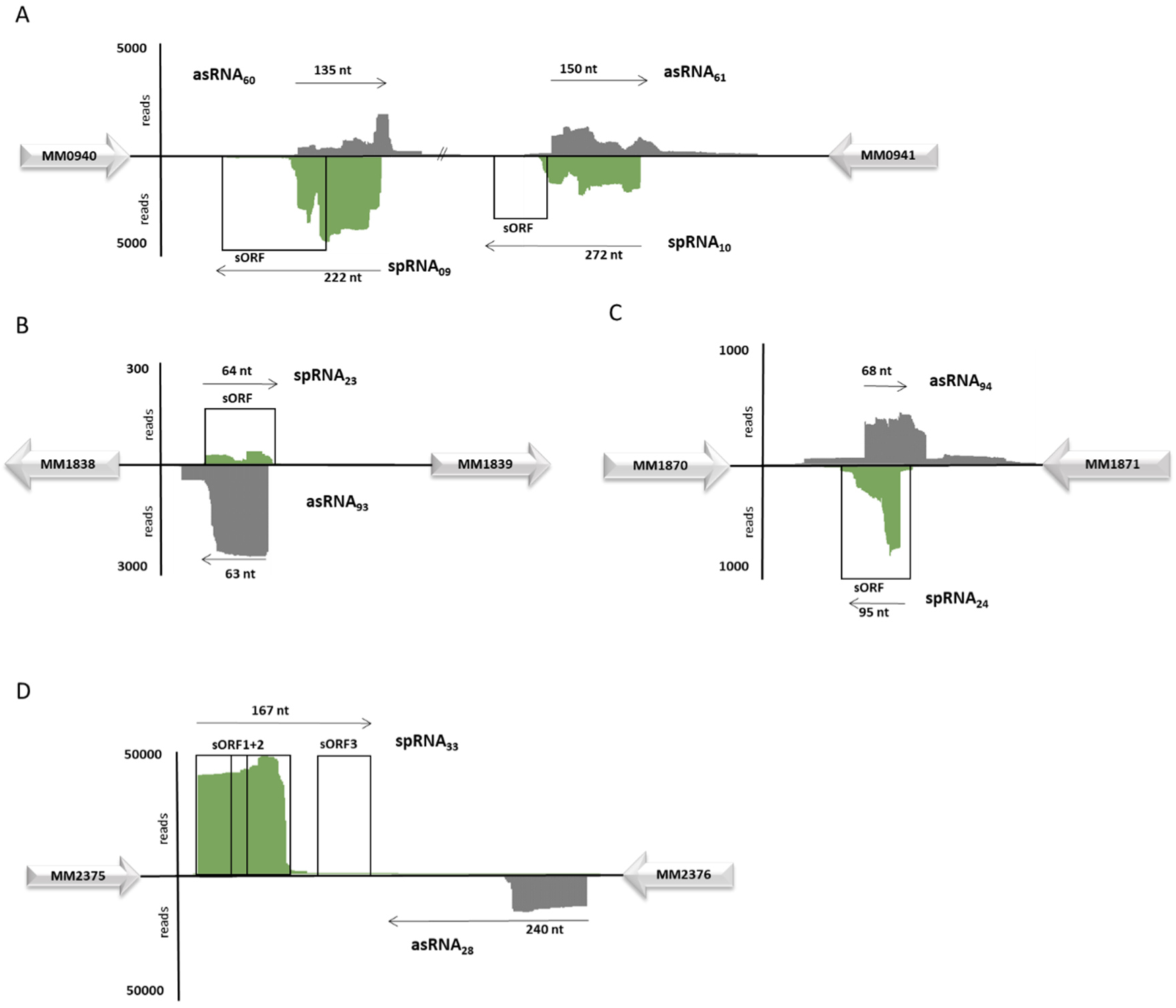
Localization in the genome and transcription of spRNAs and asRNAs in *M. mazei*. Cells were grown under nitrogen sufficient conditions, RNA extraction and RNAseq were performed and described by Nickel et al. (Nickel et al., 2013). Sequencing reads are mapped to the *M. mazei* genome (asRNA are shown in grey, spRNA are shown in green). The y axes indicate the relative score of read numbers per nucleotide. The x-axes show the length of the respective transcripts. **A**, spRNA09/asRNA60 and spRNA10/asRNA61; **B**, spRNA23/asRNA93; **C**, spRNA24/asRNA94; **D**, spRNA33/asRNA28. Predicted sORFs are depicted with boxes. Flanking genes are indicated, MM0940, putative flavoprotein; MM0941, adenylosuccinate lyase; MM1838, ferredoxin; MM1871, vanillate decarboxylase protein; MM1839, MM1870, MM2375 and MM2376 are conserved proteins with unknown function.

Sequence conservation analysis revealed that all five spRNAs are highly conserved among *Methanosarcina* species, with the exception of spRNA24, which is conserved only in a subset of *M. mazei* strains. Multiple sequence alignments and secondary structure predictions (Fig. 2) were performed using LocaRNA (Raden et al., 2018; Will, 2024; Will et al., 2012; Will et al., 2007), based on homologous sequences identified via BLASTn. Predicted start and stop codons of the sORFs are indicated in Fig. 2, alongside experimentally determined transcriptional start sites (+1) from Jäger et al. (Jäger et al., 2009), indicating that spRNA23, spRNA24, and spRNA33 are most likely transcribed as leaderless mRNAs. Predicted amino acid sequences of the corresponding small proteins show a high degree of conservation across species (Fig. S1).

**Figure 2:**
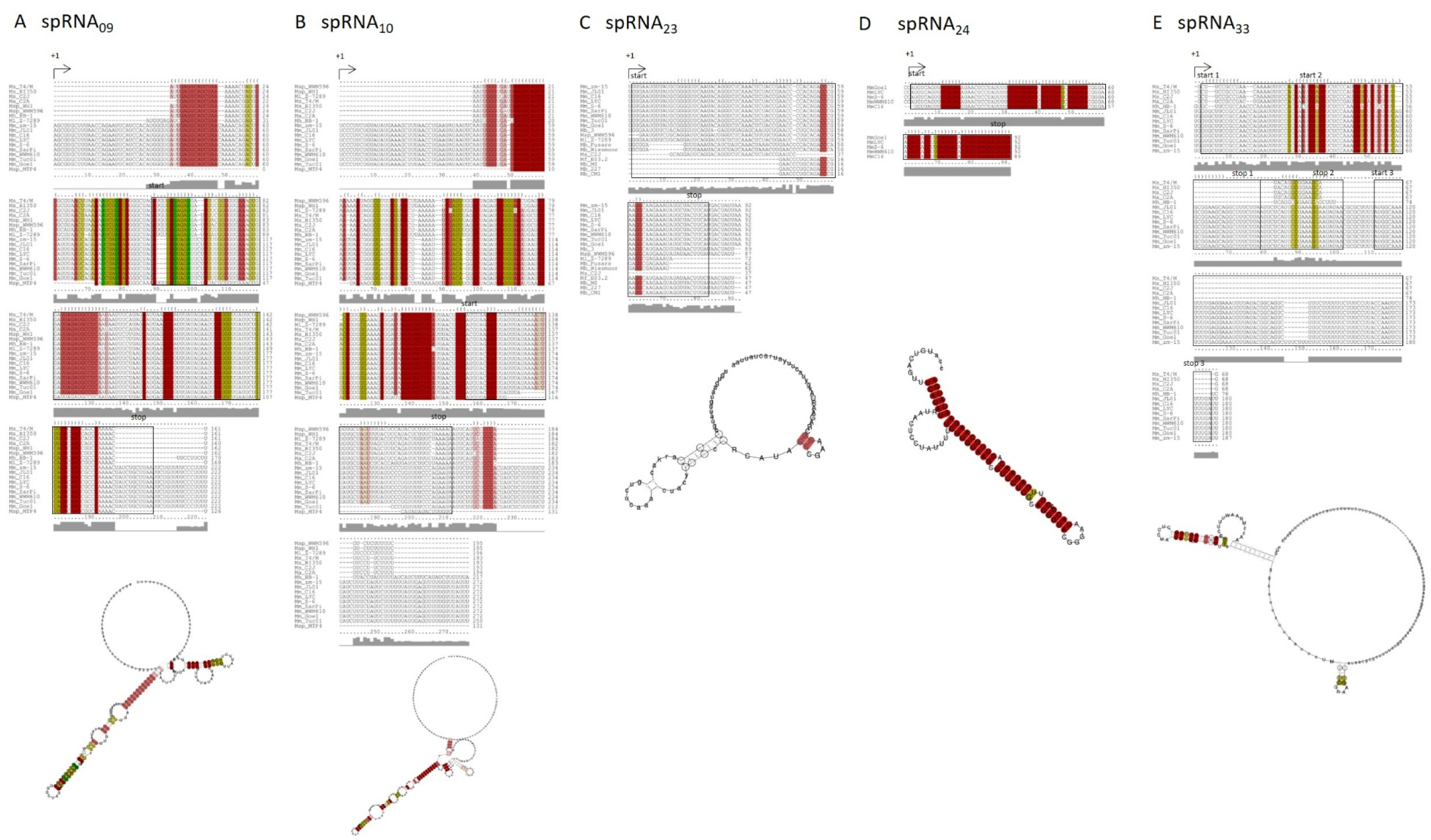
Sequence and structure conservation of spRNAs in *Methanosarcina* species. Multiple secondary structure alignment of spRNA homologues in Methanosarcina strains ((**A**) spRNA09, (**B**) spRNA10, (**C**) spRNA23, (**D**) spRNA24, (**E**) spRNA33) performed with LocaRNA (Will et al., 2007; Will et al., 2012; Raden et al., 2018). The start and stop codons of the predicted sORFs are indicated as well as the verified transcriptional start sites (+1). Below the alignments the predicted RNA structure are shown. Mm, *Methanosarcina mazei* strains zm-15, JL01, C16, LYC, S-6, SarPi, WWM610, Tuc01, Goe1; Mh: *Methanosarcina horonobensis* strain HB-1; Ma, *Methanosarcina acetivorans* strain C2A; Msp, *Methanosarcina sp.* MTP4, WH1, WWM596; Ml, *Methanosarcina alacustris* strain Z-7289; Mb, *Methanosarcina barkeri* strains Fusaro, Wiesmoor, MS, 227, CM1; Ms, *Methanosarcina siciliae* strains T4/M, HI350, C2J; Mf, *Methanosarcina flavescens* strain E03.2.

Analysis of secondary structure elements reveals that spRNA09 and spRNA10 exhibit a balanced distribution of stem and loop structures (Fig. 2A and 2B). In contrast, spRNA23 and spRNA33 are characterized by predominantly not structured (Fig. 2C and 2E, lower panel), while spRNA24 contains a conserved stem structure as its primary feature (Fig. 2D, lower panel).

### 3.2 Stress-Dependent Differential Expression of spRNA/asRNA Pairs in *M. mazei*

To investigate the expression patterns of the selected spRNA/asRNA pairs by Northern blot analysis, total RNA was extracted from *M. mazei* Gö1 cultures grown under various stress conditions. Transcript levels normalized to the 5S rRNA were analyzed during exponential and stationary phases, as well as during exponential growth under specific stress conditions (oxidative stress, growth at 30 °C and 42 °C, and 500 mM NaCl). Fold-changes in transcript abundance under stress conditions were calculated relative to the expression levels observed during the exponential phase under nitrogen-sufficient conditions (Fig. 3, upper two panels; including representative northern blots). Based on those data, the ratios of sense to antisense transcript levels were calculated for each spRNA/asRNA pair (Fig. 3, lower panel). For the spRNA23/asRNA93 pair, the sense-to-antisense ratio was strongly shifted in favor of spRNA23 under oxidative stress (8-fold), high temperature at 42 °C (14-fold), and high salinity (13-fold), as shown in Figure 3C lower panel. The spRNA24/asRNA94 pair displayed more moderate regulation, with the ratio shifted toward spRNA24 by 1.5-fold at 30 °C, 2.6-fold at 500 mM NaCl, and toward asRNA94 by 2.0-fold at 42 °C (Fig. 3D, lower panel). Due to the pronounced and condition-specific regulation observed in these two pairs, spRNA23/asRNA93 and spRNA24/asRNA94 were selected for further characterization.

**Figure 3:**
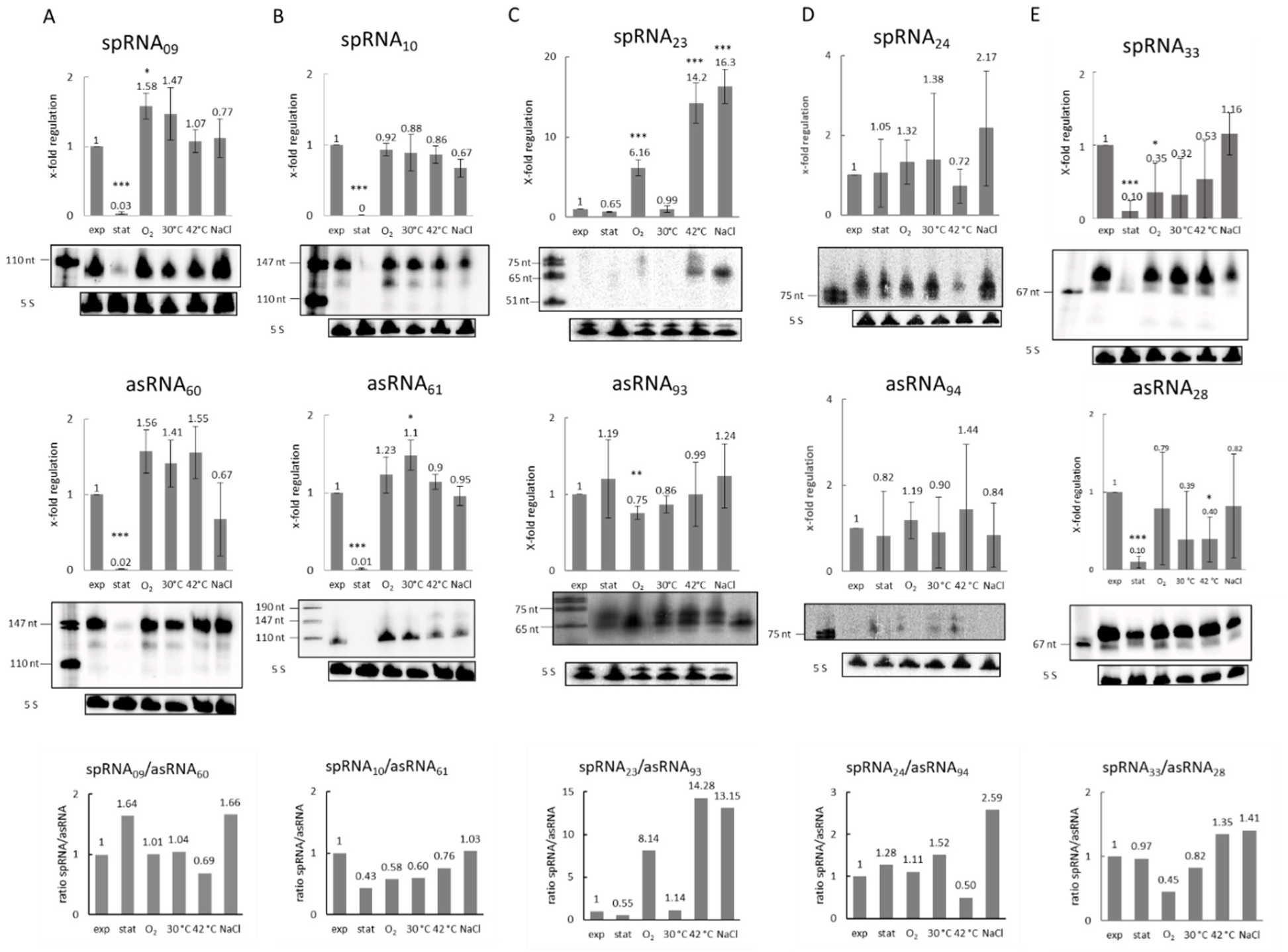
Transcript level of spRNA and asRNA in *M. mazei* grown under various stress conditions. Total RNA was isolated from *M. mazei* wild type grown under different conditions and subjected to northern blot analysis. The x fold-induction of transcript levels in exponential phase (exp), stationary phase (stat), as well as oxygen-stress (O_2_), 30 °C, 42 °C and 500 mM NaCl stress during exponential phase under N-sufficiency is calculated with respect to the transcript level in exponential phase under standard conditions after normalization to the respective 5S rRNA transcripts. The depicted standard deviation is based on three independent biological experiments. Exemplarily one original northern blot showing stable sRNA transcripts is shown for one biological replicate; the bottom panel represents the 5S rRNA transcripts of the respective RNA preparation. Some of the northern blot membranes were stripped and used with several different labeled probes, resulting in the same respective 5S rRNA control. The used labeled probes were directed against (**A**) spRNA09 and asRNA60, (**B**) spRNA10 and asRNA61, (**C**) spRNA23 and asRNA93, (**D**) spRNA24 and asRNA94, (**E**) spRNA33 and asRNA28 and are summarized in Table S1. Asterisks indicate for statistical significance (p < 0.001 = ***, p < 0.01 = **, p < 0.05 = *). Lower panel: Based on the northern blot analyses the relative x-fold regulation under each growth condition of the spRNA in relation to the corresponding asRNA was calculated, by setting the ratio of both RNAs under standard growth in the exponential phase to 1, as standard transcription pattern.

### 3.3 Transmembrane Domain Prediction Indicates Membrane Localization of sP24

Small proteins particularly those encoded by sRNAs, often exert their function through interactions with membranes or membrane-associated complexes. To gain further insight into the potential functions of the small proteins encoded by spRNA23 and spRNA24, *in silico* predictions of transmembrane domains were performed. This predicts that small protein 24 (sP24) has a transmembrane domain (Fig. 4A, using the online tools Protter and TMHMM (Krogh et al., 2001; Omasits et al., 2014)), supporting its classification as a potential transmembrane protein. The predicted domain spans the central region of the peptide (Fig. 4B), suggesting a possible role in membrane modulation. In contrast, small protein 23 did not show any predicted transmembrane regions (Fig. 4C), indicating that it is likely a soluble cytoplasmic protein.

**Figure 4.**
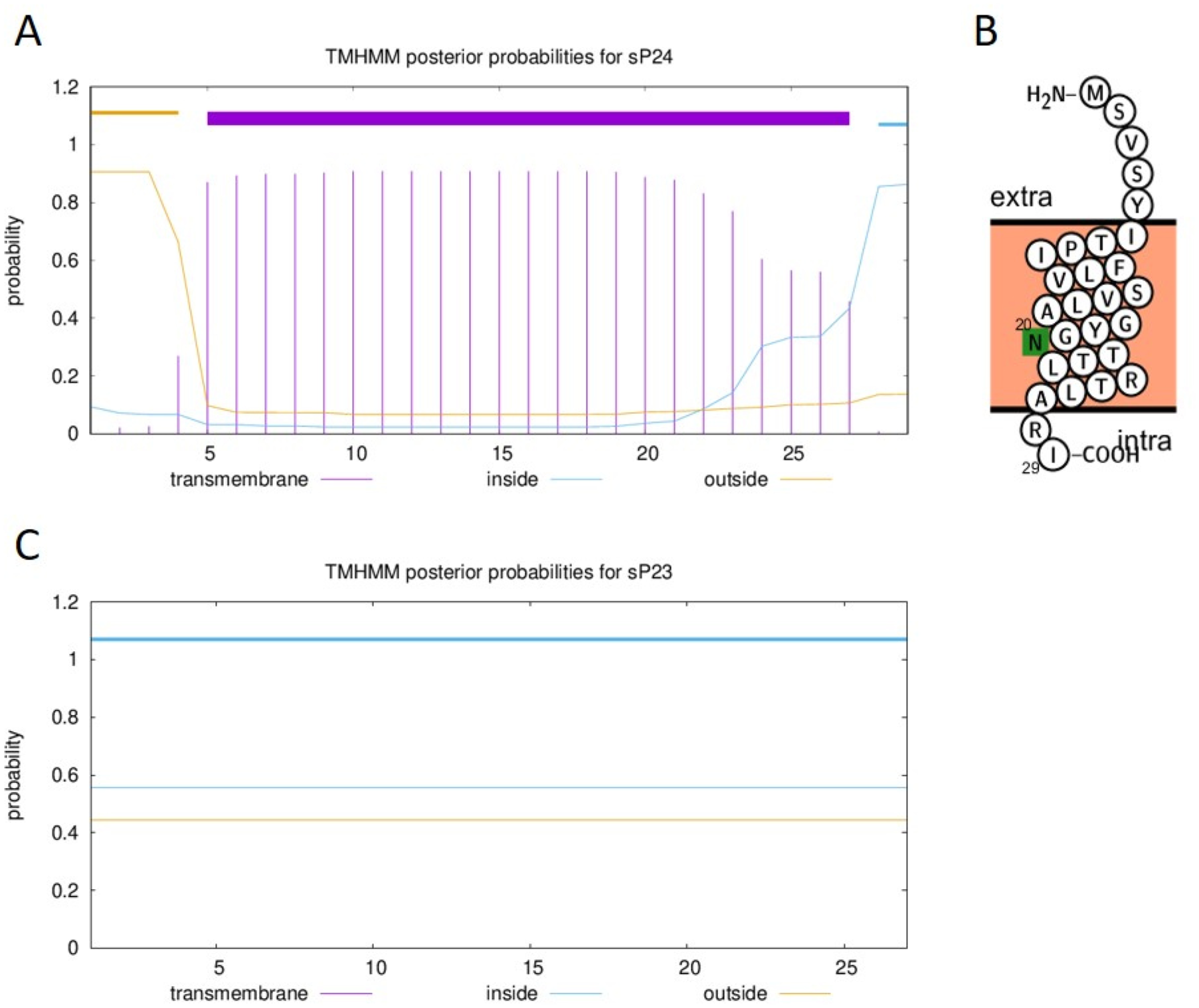
*In silico* transmembrane domain prediction of small proteins sP24 and sP23. (**A**) TMHMM (Krogh et al., 2001) posterior probability plot for sP24 shows a strong transmembrane domain prediction (purple bar) with high confidence (posterior probability > 0.9), suggesting a membrane-integrated protein topology. (**B**) Schematic topology model of sP24 generated with Protter (Omasits et al., 2014), indicating a single transmembrane helix, an extracellular N-terminus, and an intracellular C-terminus. An N-glycosylation motif is marked in green. (**C**) TMHMM (Krogh et al., 2001) posterior probability plot for sP23 reveals no predicted transmembrane domains, with uniform probabilities indicating a cytoplasmic localization.

To further validate the predicted membrane localization of sP24, we employed an *in vitro* cell-free expression system with membrane-mimicking environments as described previously (Cho et al., 2022; Bhattacharya et al., 2020; Jiang et al., 2024). In this system, sP24 was efficiently synthesized and successfully integrated into synthetic lipid bilayers, confirming its capacity to associate with membrane structures *in vitro* (Figure 5). Together, these findings provide strong indications for the membrane localization and potential structural role of sP24 during osmotic stress adaptation in *M. mazei*.

**Figure 5.**
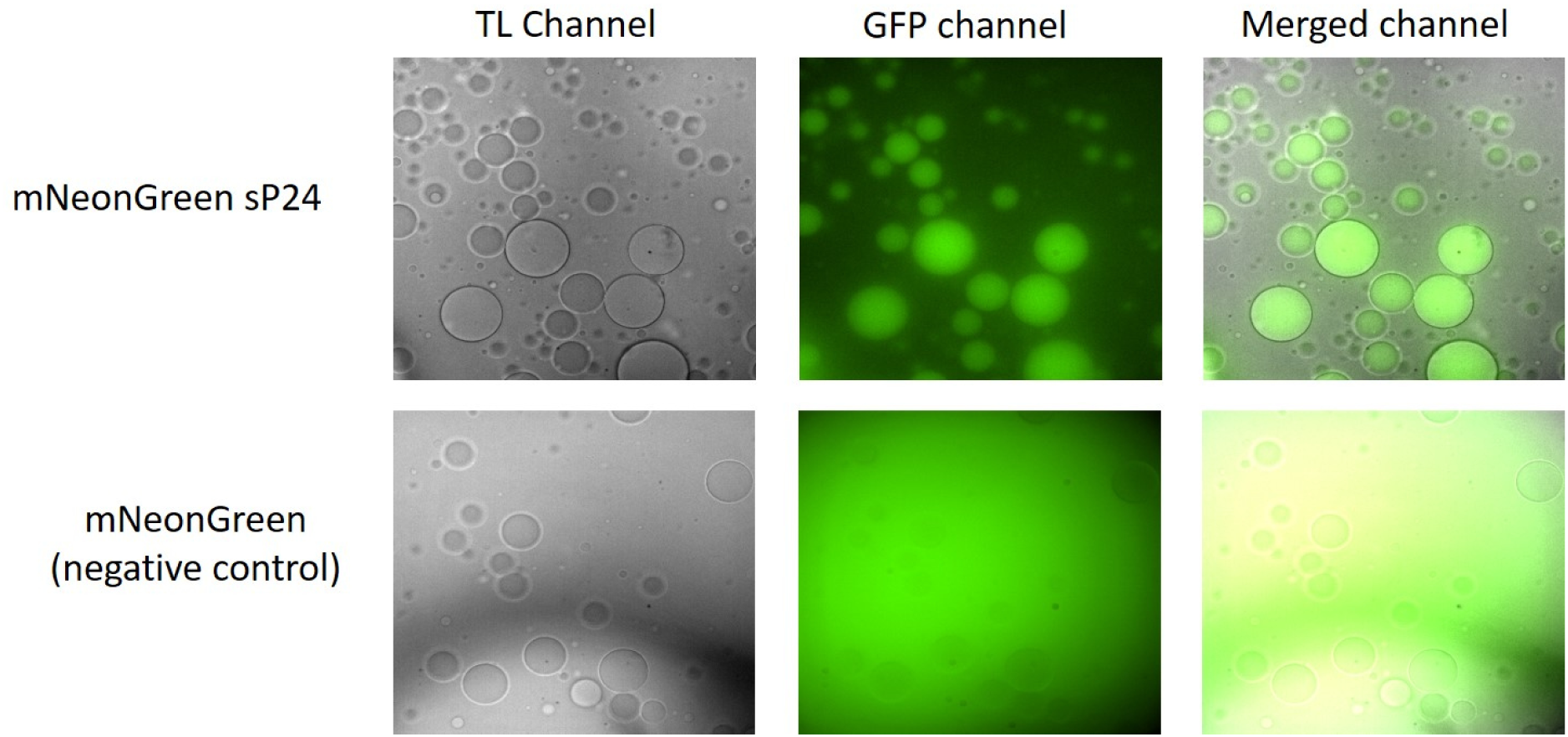
Cell-free expression of sP24 confirms membrane localization using lipid sponge droplets. The sORF24 was fused to *mneonGreen* after codon optimization and expressed in a cell-free system containing lipid sponge droplets as membrane mimics (see Materials and Methods for details). Top row: Fluorescence microscopy of mNeonGreen labeled sP24 shows clear enrichment of fluorescence in the lipid sponge droplets, indicating successful membrane integration. Images represent transmitted light (TL), GFP fluorescence, and merged channels. Bottom row: Control expression of unfused mNeonGreen results in diffuse fluorescence without vesicle association, indicating the absence of membrane interaction. These results provide direct experimental evidence supporting the membrane-associating nature of sP24, as predicted by *in silico* analyses, and further underline its potential role in membrane-related stress response mechanisms in *M. mazei*.

### 3.4 Impact of spRNAs and asRNAs on Growth Dynamics Under Stress Conditions

To investigate the role of selected RNAs and their encoded proteins, overexpression strains of *M. mazei* were generated for asRNA93, asRNA94, and the predicted small proteins sP23 and sP24, encoded by sORF23 and sORF24, respectively. Under standard growth conditions no significant impact on growth was obtained, except overexpression of asRNA94 led to a slightly reduced growth rate and an earlier entry into the stationary phase compared to the empty vector control (Fig. 6A). sORF23 as well as sORF24 overexpression resulted in significantly higher growth rates under high-salinity conditions (500 mM NaCl, Fig.6B right panel), indicating that they contribute to cellular adaptation under osmotic stress. In agreement, overproduction of the corresponding asRNA94 showed a pronounced growth defect, with a significantly prolonged lag phase and lower maximal turbidity (Fig. 6B, left panel). In contrast, overproduction of asRNA93 did not show a negative impact on the growth behavior. Shifting the temperature from 37 °C to 42 °C did not show a pronounced growth phenotype, neither for the overproduction of the asRNAs nor for the corresponding overexpressed sORFs (Fig. S2).

**Figure 6:**
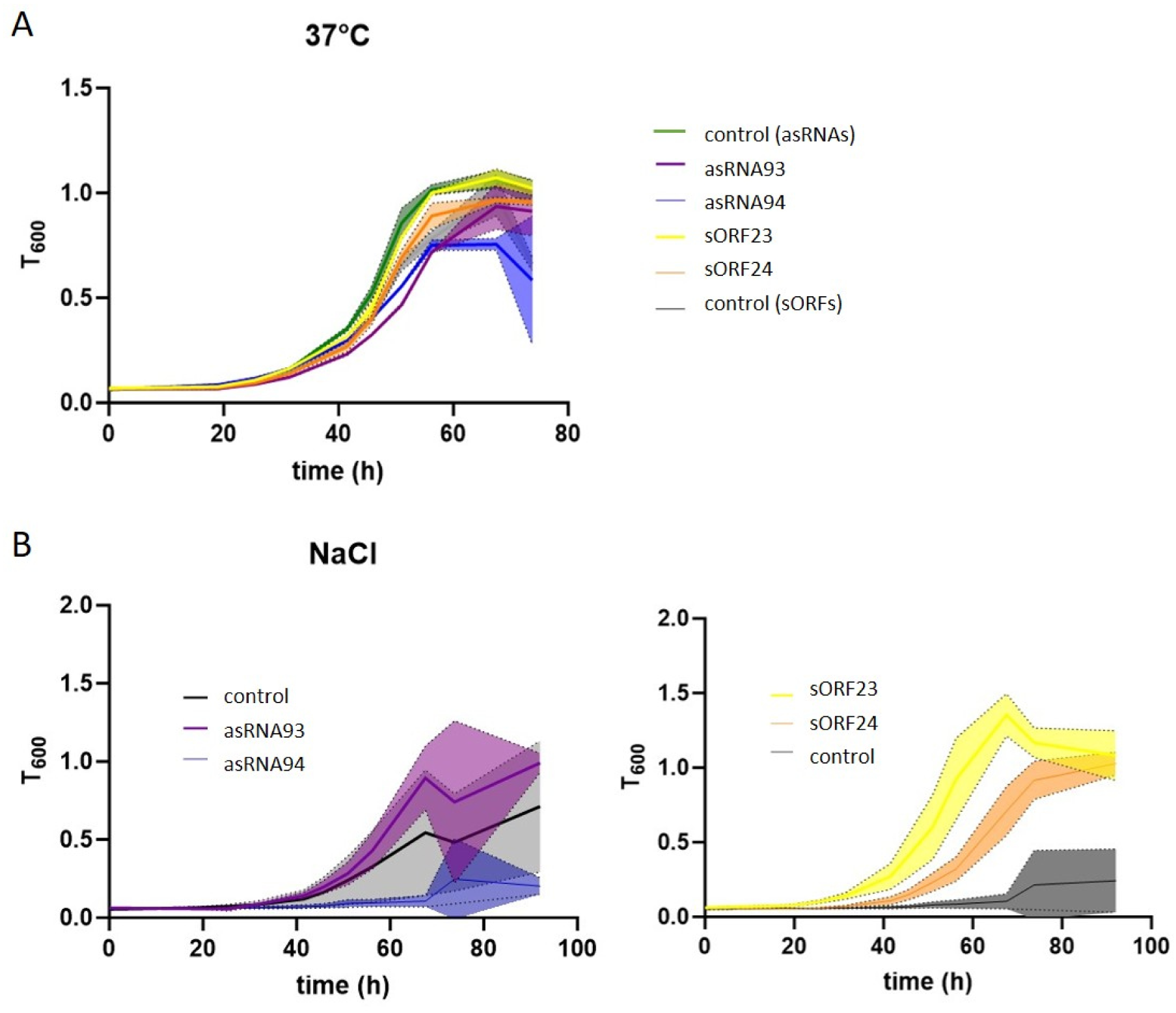
Growth analysis of *M. mazei* with additional expression of sORF23 or sORF24 and their respective asRNAs. *M. mazei* overproduction mutants, expressing asRNA93, asRNA94, sORF23 and sORF24 each under a constitutive promoter as described in Material and Methods, were cultivated in 50 ml. As a control *M. mazei* containing the empty shuttle-vectors pWM321 (control asRNAs) or pRS1595 (control sORFs) were used. (**A**) Standard growth conditions; (**B**) Cultivation under under salt stress (500 mM NaCl). The optical density at 600 nm (T_600_) was monitored. The depicted standard deviation (shadows) is based on three individually grown cultures (biological replictaes).

Overall, these results suggest a potential beneficial role for sP23 and sP24 during salt stress which may play a critical role in maintaining normal cell physiology. A dynamic balance between spRNA and corresponding asRNA in wild-type likely ensures fine-tuned regulation of translation, which adjusts in response to stress (see also Fig. 3C,D).

### 3.5 sP24 Supports Cell Size Maintenance During Osmotic Stress in *M. mazei*

Scanning electron microscopy was used to examine cell morphology and transmission electron microscopy (TEM) was performed to examine cell size in *M. mazei* cells of the empty vector control and of the two strains overexpressing the small proteins sORF23 or sORF24. Cultures were analyzed under both, standard growth conditions and osmotic stress induced by high-salt (500 mM NaCl) medium (Fig. 7). Under standard conditions, cells of all strains had a rather irregular shape and a folded cell surface as observed with SEM. Under salt stress, cell shape became angular and cell surface became smooth in all three strains (Fig. 7A). However, cells overproducing sORF24 appeared larger than the other two strains and to validate this observation, cell size was estimated using TEM images of stained whole mount cells (Fig. 7A). No significant differences in cell size were observed between strains under standard conditions. Under salt stress, size of wild-type cells showed a 28% decrease compared to the standard conditions, while cells overproducing sORF23 exhibited an even greater size reduction of 35%. Notably, overproduction of sORF24 significantly weakened the stress-induced shrinkage, with only a 15% reduction in cell size. In consequence, cells overproducing sORF24 were significantly larger than empty vector control cells or cells overproducing sORF23 (p < 0.01) (Fig. 7C). These findings suggest that overproduction of either sORF does not affect cell morphology under standard conditions. However, sORF24 contributes to cellular resilience during osmotic stress, likely by stabilizing membrane integrity or influencing structural components, consistent with its transmembrane nature. This protective effect highlights a functional role in maintaining cell size under adverse environmental conditions.

**Figure 7:**
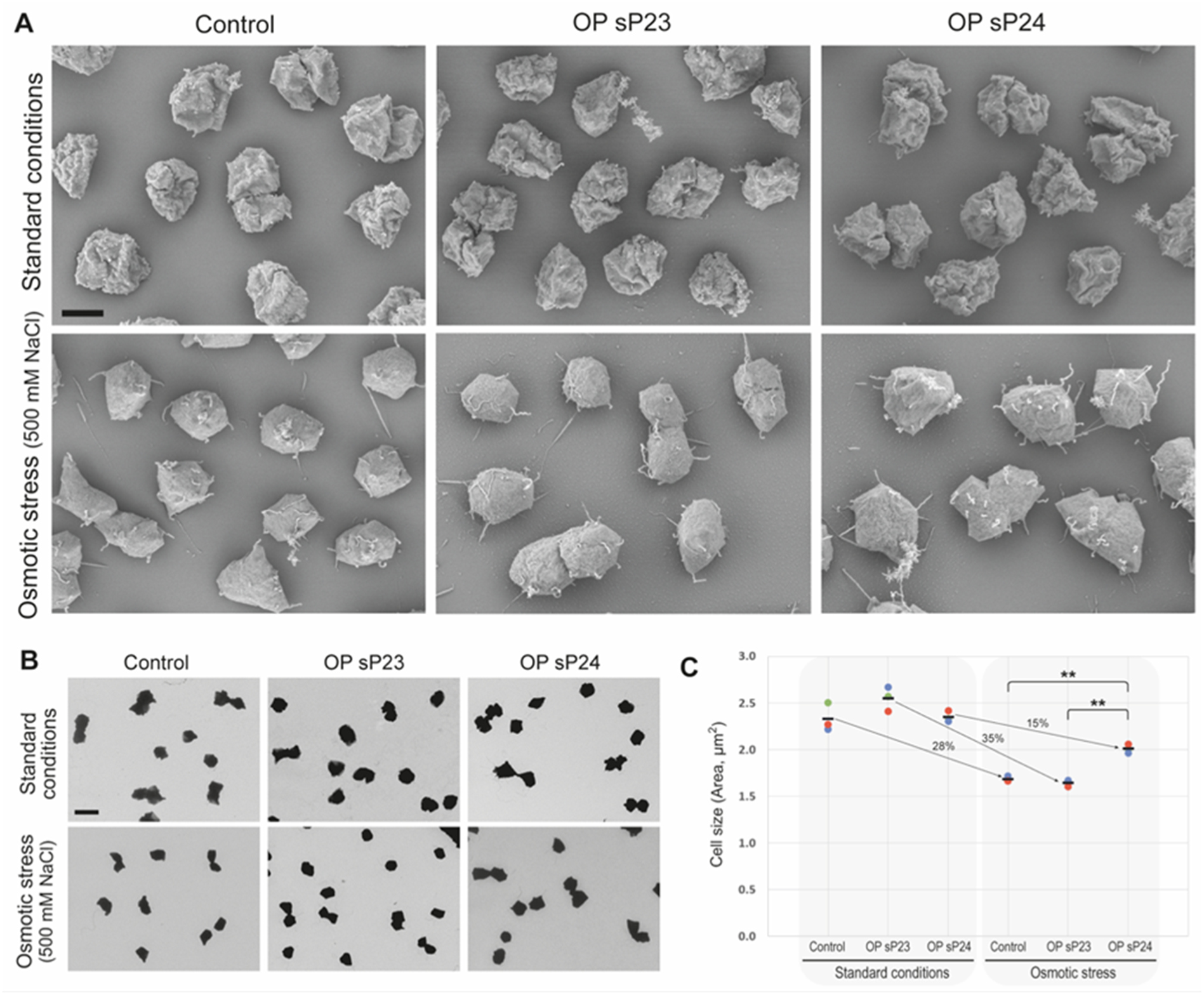
Morphological and cell size analyses under standard conditions and under osmotic stress. (A) Cell shape visualized by scanning electron microscopy. Scalebar, 1 μm. (B,C) Cell size analysis based on transmission electron micrographs of whole cells. (B) Cell size was estimated as the area of the cell projection. Scalebar, 2.5 μm. (C) In the graph, coloured points represent mean values (n=600) of three independent experiments, black lines indicate the population mean. Under standard conditions (left panel) no significant changes in size were observed. Under high salt stress, cell size decreased, but significantly less in sP24 overproducing cells. Statistical significance of differences was calculated with a post-hoc Tukey Honest Significant Difference (HSD), ** (p<0.01).

## 4 Discussion

The discovery and characterization of small proteins and their regulatory RNAs represent a growing frontier in molecular microbiology. Despite progress in bacteria, little is known about how archaea employ sRNAs, asRNAs, and the encoded small proteins to regulate cellular function, especially under environmental stress conditions. In this study, we provide evidence that *M. mazei* utilizes two spRNA/asRNA pairs, spRNA23/asRNA93 and spRNA24/asRNA94, to modulate growth in response to salt stress, most likely through their encoded small proteins, sP23 and sP24.

Post-transcriptional regulation by sRNAs and asRNAs is well established in bacteria, with classic toxin/antitoxin systems like Sok–Hok in *Escherichia coli*. This system controls the synthesis of a small hydrophobic toxin (Hok) via the antisense RNA Sok, which inhibits translation and thereby prevents cell death under non-stress conditions (Gerdes et al., 1986; Gerdes et al., 1990). Similarly, the SymR–SymE pair regulates SymE, a stress-induced ribonuclease, with the asRNA SymR acting as a repressor of SymE translation (Kawano et al., 2007; Thompson et al., 2022). These examples illustrate how antisense RNAs suppress the translation of potentially toxic small proteins. Similar strategies may be conserved in archaea. In our study, spRNA23 and spRNA24 showed stress-dependent induction (oxidative, temperature, and high osmotic conditions), but spRNA24 shows down regulation at 42°C. The corresponding asRNAs generally exhibited opposite regulation. This reciprocal expression pattern suggests that asRNAs, such as asRNA93 and asRNA94, may serve as repressors that fine-tune small protein production, preventing unnecessary or harmful expression under non-stress conditions (Fozo et al., 2010; Kawano, 2012). Overexpression of particularly asRNA94 resulted in notable growth defects (Fig. 6), supporting its predicted role as translational silencer. These findings strengthen the concept that balanced production between sense and antisense transcripts is essential for normal cell physiology and that disruption of this balance impairs growth (Edelmann & Berghoff, 2022).

The small proteins encoded by spRNA23 and spRNA24 have been shown to be biologically relevant under stress. Notably, sORF23 overexpression enhanced growth under high-salt conditions (Fig. 6B), suggesting a beneficial role, possibly in membrane stabilization or stress response signaling, which is in agreement with the significant upregulation of spRNA23 under salt stress (Fig. 3C). Similarly, sP24, predicted to be a transmembrane protein, significantly enhanced growth and weakened the typical reduction in cell size during osmotic stress (see Fig. 6B and Fig.7), strongly indicating a protective effect on membrane integrity or osmotic balance. Membrane localization of sORF24 shown by a cell-free lipid membrane integration assay (Jiang et al., 2024) (Fig. 5), strongly indicates in vitro ability to integrate into hydrophobic lipid environments, further supporting its potential membrane-associated function.

Similarly, membrane-associated localization and function have been observed for several bacterial small proteins. For instance, Prli42 in *Listeria monocytogenes* is a 31 amino acid (aa) transmembrane protein that interacts with the stressosome complex RsbRST to initiate general stress responses (Impens et al., 2017; Williams et al., 2019). In *E. coli*, the small protein YohP (27 aa) localizes to the inner membrane and is upregulated under various stress conditions, including minimal media and low oxygen (Hemm et al., 2010). These analogies strongly suggest that the small protein sP24 may interact with the cytoplasmic membrane or associated complexes to modulate stress adaptation in *M. mazei*.

Conservation analysis showed that while most identified spRNAs are conserved within the *Methanosarcina* genus, spRNA24 appears to be strain-specific, hinting at adaptation to specific environmental niches. Such genomic variability may reflect evolutionary fine-tuning of regulatory systems to match habitat-specific stressors (Steinberg & Koch, 2021).

Moreover, many previously annotated non-coding RNAs in *M. mazei* appear to encode functional small proteins acting potentially as dual function RNAs (Tufail et al., 2024). In this respect, optimized mass spectroscopy for small proteins and the application of Ribo-seq have been instrumental in uncovering such sORFs in both bacteria and archaea, including *Haloferax volcanii* and *M. mazei* (Gelsinger et al., 2020; Hadjeras et al., 2023; Tufail et al., 2024; Vazquez-Laslop et al., 2022). These tools are currently rapidly expanding our understanding of genome functionality and regulatory complexity in prokaryotes.

Overall this study adds to the growing body of work highlighting the biological importance of small RNAs and small proteins in archaea. It demonstrates that even short peptides, once overlooked due to their size, can exert significant physiological effects, particularly under stress. The interaction between sRNAs and their antisense counterparts might be central to this regulation and appears to follow principles broadly conserved across domains (Hobbs et al., 2011; Song et al., 2022). Moving forward, further characterization of sORF23 and sORF24 will benefit from membrane fractionation, subcellular localization, and interaction proteomics to uncover their precise cellular roles. In parallel, knockout or conditional expression systems could confirm their contribution to fitness under fluctuating environmental conditions and deepen our understanding of stress adaptation in archaea.

## 5 Conclusion

The coordinated expression and regulation of spRNA23/asRNA93 and spRNA24/asRNA94 in *M. mazei* are key components of the organism’s adaptive response to environmental stress. The small proteins they encode, sP23 and sP24, enhance cellular fitness under osmotic stress, with the membrane-associated sP24 particularly contributing to the preservation of cell size. Conversely, overexpression of asRNA94 interferes with this balance, leading to growth defects. Together, these findings underscore the functional significance of sRNA/asRNA pairs in archaeal stress responses and the emerging regulatory roles of small proteins encoded by non-coding RNAs.

## 7 Author contribution

RAS conceptualized and designed the study; RAS, JY, BJ, UR, DP and RG contributed to methodology and investigated the study; BJ, RG, UR and SJ visualized the study; BJ, RAS, JY, and LH wrote original draft; RAS supervised the study and was responsible for funding acquisition.

## 9 Acknowledgement

We thank the members of our laboratories for useful discussions on the experiments, as well as Claudia Kiessling for technical assistance. We acknowledge the Central Microscopy facility at Christian-Albrechts-University, Faculty of Mathematics and Natural Sciences, Dept Biology, for electron microscopic analysis.

## References

Aoyama, J. J., Raina, M., Zhong, A., & Storz, G. (2022). Dual-function Spot 42 RNA encodes a 15-amino acid protein that regulates the CRP transcription factor. Proc Natl Acad Sci U S A, 119(10), e2119866119. 10.1073/pnas.2119866119

Babski, J., Haas, K. A., Näther-Schindler, D., Pfeiffer, F., Förstner, K. U., Hammelmann, M., Hilker, R., Becker, A., Sharma, C. M., Marchfelder, A., & Soppa, J. (2016). Genome-wide identification of transcriptional start sites in the haloarchaeon Haloferax volcanii based on differential RNA-Seq (dRNA-Seq). BMC Genomics, 17(1), 629. 10.1186/s12864-016-2920-y

Babski, J., Tjaden, B., Voss, B., Jellen-Ritter, A., Marchfelder, A., Hess, W. R., & Soppa, J. (2011). Bioinformatic prediction and experimental verification of sRNAs in the haloarchaeon Haloferax volcanii. RNA Biol, 8(5), 806–816. 10.4161/rna.8.5.16039

Bernick, D. L., Karplus, K., Lui, L. M., Coker, J. K., Murphy, J. N., Chan, P. P., Cozen, A. E., & Lowe, T. M. (2012). Complete genome sequence of Pyrobaculum oguniense. Stand Genomic Sci, 6(3), 336–345. 10.4056/sigs.2645906

Bhattacharya A, Niederholtmeyer H, Podolsky KA, Bhattacharya R, Song J, Brea JR, Tsai C, Sinha SK, Devaraj NK (2020), Lipid sponge droplets as programmable synthetic organelles, Proc. Natl. Acad. Sci. U.S.A. 117 (31) 18206–18215, 10.1073/pnas.2004408117 (2020)

Buddeweg, A., Sharma, K., Urlaub, H., & Schmitz, R. A. (2018). sRNA(41) affects ribosome binding sites within polycistronic mRNAs in Methanosarcina mazei Gö1. Mol Microbiol, 107(5), 595–609. 10.1111/mmi.13900

Burton, A. T., Zeinert, R., & Storz, G. (2024). Large Roles of Small Proteins. Annu Rev Microbiol, 78(1), 1–22. 10.1146/annurev-micro-112723-083001

Cho, C. J., Niederholtmeyer, H., Seo, H., Bhattacharya, A., Devaraj, N. K. (2022) Functionalizing Lipid Sponge Droplets with DNA**. ChemSystemsChem 2022, *4*, e202100045 10.1002/syst.202100045.

Chihara, K., Gerovac, M., Hör, J., & Vogel, J. (2022). Global profiling of the RNA and protein complexes of Escherichia coli by size exclusion chromatography followed by RNA sequencing and mass spectrometry (SEC-seq). Rna, 29(1), 123–139. 10.1261/rna.079439.122

Deppenmeier, U., Blaut, M., Mahlmann, A., & Gottschalk, G. (1990). Reduced coenzyme F_420_: heterodisulfide oxidoreductase, a proton-translocating redox system in methanogenic bacteria. Proc Natl Acad Sci U S A, 87(23), 9449–9453. http://www.ncbi.nlm.nih.gov/entrez/query.fcgi?cmd=Retrieve&db=PubMed&dopt=Citation&list_uids=11607121

Deppenmeier, U., Johann, A., Hartsch, T., Merkl, R., Schmitz, R. A., Martinez-Arias, R., Henne, A., Wiezer, A., Baumer, S., Jacobi, C., Bruggemann, H., Lienard, T., Christmann, A., Bomeke, M., Steckel, S., Bhattacharyya, A., Lykidis, A., Overbeek, R., Klenk, H. P., . . . Gottschalk, G. (2002). The genome of *Methanosarcina mazei*: evidence for lateral gene transfer between bacteria and archaea. J Mol Microbiol Biotechnol, 4(4), 453–461. http://www.ncbi.nlm.nih.gov/entrez/query.fcgi?cmd=Retrieve&db=PubMed&dopt=Citation&list_uids=12125824

Deppenmeier, U., Müller, V., & Gottschalk, G. (1996). Pathways of energy conservation in methanogenic archaea. Arch Microbiol, 165, 149–163.

Edelmann, D., & Berghoff, B. A. (2022). A Shift in Perspective: A Role for the Type I Toxin TisB as Persistence-Stabilizing Factor. Front Microbiol, 13, 871699. 10.3389/fmicb.2022.871699

Ehlers, C., Jäger, D., & Schmitz, R. A. (2011). Establishing a markerless genetic exchange system for *Methanosarcina mazei* strain Gö1 for constructing chromosomal mutants of small RNA genes. Archaea, 2011, 439608. http://www.ncbi.nlm.nih.gov/entrez/query.fcgi?cmd=Retrieve&db=PubMed&dopt=Citation&list_uids=21941461

Ehlers, C., Veit, K., Gottschalk, G., & Schmitz, R. A. (2002). Functional organization of a single nif cluster in the mesophilic archaeon Methanosarcina mazei strain Gö1. Archaea, 1(2), 143–150. 10.1155/2002/362813

Ehlers, C., Weidenbach, K., Veit, K., Deppenmeier, U., Metcalf, W. W., & Schmitz, R. A. (2005). Development of genetic methods and construction of a chromosomal glnK1 mutant in Methanosarcina mazei strain Gö1. Mol Genet Genomics, 273(4), 290–298. 10.1007/s00438-005-1128-7

Ferry, J. G. (1999). Enzymology of one-carbon metabolism in methanogenic pathways. FEMS Microbiol Rev, 23(1), 13–38. http://www.ncbi.nlm.nih.gov/entrez/query.fcgi?cmd=Retrieve&db=PubMed&dopt=Citation&list_uids=10077852

Fozo, E. M., Makarova, K. S., Shabalina, S. A., Yutin, N., Koonin, E. V., & Storz, G. (2010). Abundance of type I toxin-antitoxin systems in bacteria: searches for new candidates and discovery of novel families. Nucleic Acids Res, 38(11), 3743–3759. 10.1093/nar/gkq054

Gehlert, F. O., Nickel, L., Vakirlis, N., Hammerschmidt, K., Vargas Gebauer, H. I., Kießling, C., Kupczok, A., & Schmitz, R. A. (2023). Active in vivo translocation of the Methanosarcina mazei Gö1 Casposon. Nucleic Acids Res, 51(13), 6927–6943. 10.1093/nar/gkad474

Gelsinger, D. R., Dallon, E., Reddy, R., Mohammad, F., Buskirk, A. R., & DiRuggiero, J. (2020). Ribosome profiling in archaea reveals leaderless translation, novel translational initiation sites, and ribosome pausing at single codon resolution. Nucleic Acids Res, 48(10), 5201–5216. 10.1093/nar/gkaa304

Gerdes, K., Rasmussen, P. B., & Molin, S. (1986). Unique type of plasmid maintenance function: postsegregational killing of plasmid-free cells. Proc Natl Acad Sci U S A, 83(10), 3116–3120. 10.1073/pnas.83.10.3116

Gerdes, K., Thisted, T., & Martinussen, J. (1990). Mechanism of post-segregational killing by the hok/sok system of plasmid R1: sok antisense RNA regulates formation of a hok mRNA species correlated with killing of plasmid-free cells. Mol Microbiol, 4(11), 1807–1818. 10.1111/j.1365-2958.1990.tb02029.x

Ghandour, R., & Papenfort, K. (2023). Small regulatory RNAs in Vibrio cholerae. Microlife, 4, uqad030. 10.1093/femsml/uqad030

Gimpel, M., & Brantl, S. (2017). Dual-function small regulatory RNAs in bacteria. Mol Microbiol, 103(3), 387–397. 10.1111/mmi.13558

Guss, A. M., Rother, M., Zhang, J. K., Kulkarni, G., & Metcalf, W. W. (2008). New methods for tightly regulated gene expression and highly efficient chromosomal integration of cloned genes for *Methanosarcina* species. Archaea, 2(3), 193–203. http://www.ncbi.nlm.nih.gov/entrez/query.fcgi?cmd=Retrieve&db=PubMed&dopt=Citation&list_uids=19054746

Hadjeras, L., Bartel, J., Maier, L. K., Maaß, S., Vogel, V., Svensson, S. L., Eggenhofer, F., Gelhausen, R., Müller, T., Alkhnbashi, O. S., Backofen, R., Becher, D., Sharma, C. M., & Marchfelder, A. (2023). Revealing the small proteome of Haloferax volcanii by combining ribosome profiling and small-protein optimized mass spectrometry. Microlife, 4, uqad001. 10.1093/femsml/uqad001

Hanahan, D. (1983). Studies on transformation of *Escherichia coli* with plasmids. J Mol Biol, 166(4), 557–580. http://www.ncbi.nlm.nih.gov/entrez/query.fcgi?cmd=Retrieve&db=PubMed&dopt=Citation&list_uids=6345791

Hemm, M. R., Paul, B. J., Miranda-Ríos, J., Zhang, A., Soltanzad, N., & Storz, G. (2010). Small stress response proteins in Escherichia coli: proteins missed by classical proteomic studies. J Bacteriol, 192(1), 46–58. 10.1128/jb.00872-09

Hemm, M. R., Weaver, J., & Storz, G. (2020). Escherichia coli Small Proteome. EcoSal Plus, 9(1). 10.1128/ecosalplus.ESP-0031-2019

Heyer, R., Dorr, M., Jellen-Ritter, A., Spath, B., Babski, J., Jaschinski, K., Soppa, J., & Marchfelder, A. (2012). High throughput sequencing reveals a plethora of small RNAs including tRNA derived fragments in *Haloferax volcanii*. RNA Biol, 9(7), 1011–1018. http://www.ncbi.nlm.nih.gov/entrez/query.fcgi?cmd=Retrieve&db=PubMed&dopt=Citation&list_uids=22767255

Hippe, H., Caspari, D., Fiebig, K., & Gottschalk, G. (1979). Utilization of trimethylamine and other N-methyl compounds for growth and methane formation by *Methanosarcina barkeri*. Proc Natl Acad Sci U S A, 76(1), 494–498. http://www.ncbi.nlm.nih.gov/entrez/query.fcgi?cmd=Retrieve&db=PubMed&dopt=Citation&list_uids=284366

Hobbs, E. C., Fontaine, F., Yin, X., & Storz, G. (2011). An expanding universe of small proteins. Curr Opin Microbiol, 14(2), 167–173. 10.1016/j.mib.2011.01.007

Hook, S. E., Wright, A. D., & McBride, B. W. (2010). Methanogens: methane producers of the rumen and mitigation strategies. Archaea, 2010, 945785. 10.1155/2010/945785

Hör, J., Di Giorgio, S., Gerovac, M., Venturini, E., Förstner, K. U., & Vogel, J. (2020). Grad-seq shines light on unrecognized RNA and protein complexes in the model bacterium Escherichia coli. Nucleic Acids Res, 48(16), 9301–9319. 10.1093/nar/gkaa676

Impens, F., Rolhion, N., Radoshevich, L., Bécavin, C., Duval, M., Mellin, J., García Del Portillo, F., Pucciarelli, M. G., Williams, A. H., & Cossart, P. (2017). N-terminomics identifies Prli42 as a membrane miniprotein conserved in Firmicutes and critical for stressosome activation in Listeria monocytogenes. Nat Microbiol, 2, 17005. 10.1038/nmicrobiol.2017.5

Inoue, H., Nojima, H., & Okayama, H. (1990). High efficiency transformation of *Escherichia coli* with plasmids. Gene, 96(1), 23–28. http://www.ncbi.nlm.nih.gov/entrez/query.fcgi?cmd=Retrieve&db=PubMed&dopt=Citation&list_uids=2265755

Jäger, D., Pernitzsch, S. R., Richter, A. S., Backofen, R., Sharma, C. M., & Schmitz, R. A. (2012). An archaeal sRNA targeting cis- and trans-encoded mRNAs via two distinct domains. Nucleic Acids Res, 40(21), 10964–10979. 10.1093/nar/gks847

Jäger, D., Sharma, C. M., Thomsen, J., Ehlers, C., Vogel, J., & Schmitz, R. A. (2009). Deep sequencing analysis of the *Methanosarcina mazei* Gö1 transcriptome in response to nitrogen availability. Proc Natl Acad Sci U S A, 106(51), 21878–21882. http://www.ncbi.nlm.nih.gov/entrez/query.fcgi?cmd=Retrieve&db=PubMed&dopt=Citation&list_uids=19996181

Jaschinski, K., Babski, J., Lehr, M., Burmester, A., Benz, J., Heyer, R., Dörr, M., Marchfelder, A., & Soppa, J. (2014). Generation and phenotyping of a collection of sRNA gene deletion mutants of the haloarchaeon Haloferax volcanii. PLoS One, 9(3), e90763. 10.1371/journal.pone.0090763

Jiang, S., Çelen, G., Glatter, T., Niederholtmeyer, H., & Yuan, J. (2024). A cell-free system for functional studies of small membrane proteins. J Biol Chem, 300(11), 107850. 10.1016/j.jbc.2024.107850

Kawano, M. (2012). Divergently overlapping cis-encoded antisense RNA regulating toxin-antitoxin systems from E. coli: hok/sok, ldr/rdl, symE/symR. RNA Biol, 9(12), 1520–1527. 10.4161/rna.22757

Kawano, M., Aravind, L., & Storz, G. (2007). An antisense RNA controls synthesis of an SOS-induced toxin evolved from an antitoxin. Mol Microbiol, 64(3), 738–754. 10.1111/j.1365-2958.2007.05688.x

Krogh, A., Larsson, B., von Heijne, G., & Sonnhammer, E. L. (2001). Predicting transmembrane protein topology with a hidden Markov model: application to complete genomes. J Mol Biol, 305(3), 567–580. 10.1006/jmbi.2000.4315

Liu, Y. (2010). Taxonomy of Methanogens. In K. N. Timmis (Ed.), Handbook of Hydrocarbon and Lipid Microbiology (pp. 547–558). Springer Berlin Heidelberg. 10.1007/978-3-540-77587-4_42

Märtens, B., Manoharadas, S., Hasenöhrl, D., Manica, A., & Bläsi, U. (2013). Antisense regulation by transposon-derived RNAs in the hyperthermophilic archaeon Sulfolobus solfataricus. EMBO Rep, 14(6), 527–533. 10.1038/embor.2013.47

Miller, V. L., & Mekalanos, J. J. (1988). A novel suicide vector and its use in construction of insertion mutations: osmoregulation of outer membrane proteins and virulence determinants in *Vibrio cholerae* requires *toxR*. J Bacteriol, 170(6), 2575–2583. http://www.ncbi.nlm.nih.gov/entrez/query.fcgi?cmd=Retrieve&db=PubMed&dopt=Citation&list_uids=2836362

Nickel, L., Weidenbach, K., Jager, D., Backofen, R., Lange, S. J., Heidrich, N., & Schmitz, R. A. (2013). Two CRISPR-Cas systems in *Methanosarcina mazei* strain Go1 display common processing features despite belonging to different types I and III. RNA Biol, 10(5), 779–791. 10.4161/rna.23928

Omasits, U., Ahrens, C. H., Müller, S., & Wollscheid, B. (2014). Protter: interactive protein feature visualization and integration with experimental proteomic data. Bioinformatics, 30(6), 884–886. 10.1093/bioinformatics/btt607

Papenfort, K., & Melamed, S. (2023). Small RNAs, Large Networks: Posttranscriptional Regulons in Gram-Negative Bacteria. Annu Rev Microbiol, 77, 23–43. 10.1146/annurev-micro-041320-025836

Papenfort, K., & Vogel, J. (2010). Regulatory RNA in bacterial pathogens. Cell Host Microbe, 8(1), 116–127. 10.1016/j.chom.2010.06.008

Patraquim, P., Mumtaz, M. A. S., Pueyo, J. I., Aspden, J. L., & Couso, J. P. (2020). Developmental regulation of canonical and small ORF translation from mRNAs. Genome Biol, 21(1), 128. 10.1186/s13059-020-02011-5

Prasse, D., Förstner, K. U., Jäger, D., Backofen, R., & Schmitz, R. A. (2017). sRNA(154) a newly identified regulator of nitrogen fixation in Methanosarcina mazei strain Gö1. RNA Biol, 14(11), 1544–1558. 10.1080/15476286.2017.1306170

Prasse, D., & Schmitz, R. A. (2018). Small RNAs Involved in Regulation of Nitrogen Metabolism. Microbiol Spectr, 6(4). 10.1128/microbiolspec.RWR-0018-2018

Raden, M., Ali, S. M., Alkhnbashi, O. S., Busch, A., Costa, F., Davis, J. A., Eggenhofer, F., Gelhausen, R., Georg, J., Heyne, S., Hiller, M., Kundu, K., Kleinkauf, R., Lott, S. C., Mohamed, M. M., Mattheis, A., Miladi, M., Richter, A. S., Will, S., . . . Backofen, R. (2018). Freiburg RNA tools: a central online resource for RNA-focused research and teaching. Nucleic Acids Res, 46(W1), W25–w29. 10.1093/nar/gky329

Raina, M., King, A., Bianco, C., & Vanderpool, C. K. (2018). Dual-Function RNAs. Microbiol Spectr, 6(5). 10.1128/microbiolspec.RWR-0032-2018

Ramamurthi, K. S., & Storz, G. (2014). The small protein floodgates are opening; now the functional analysis begins. BMC Biol, 12, 96. 10.1186/s12915-014-0096-y

Schindelin J, Arganda-Carreras I, Frise E, Kaynig V, Longair M, Pietzsch T, Preibisch S, Rueden C, Saalfeld S, Schmid B, Tinevez JY, White DJ, Hartenstein V, Eliceiri K, Tomancak P, Cardona A. (2012). Fiji: an open-source platform for biological-image analysis. Nat Methods. 9(7):676–82. doi: 10.1038/nmeth.2019.

Schlesinger, D., & Elsässer, S. J. (2022). Revisiting sORFs: overcoming challenges to identify and characterize functional microproteins. Febs j, 289(1), 53–74. 10.1111/febs.15769

Song, K., Baumgartner, D., Hagemann, M., Muro-Pastor, A. M., Maaß, S., Becher, D., & Hess, W. R. (2022). AtpΘ is an inhibitor of F(0)F(1) ATP synthase to arrest ATP hydrolysis during low-energy conditions in cyanobacteria. Curr Biol, 32(1), 136–148.e135. 10.1016/j.cub.2021.10.051

Steinberg, R., & Koch, H. G. (2021). The largely unexplored biology of small proteins in pro- and eukaryotes. Febs j, 288(24), 7002–7024. 10.1111/febs.15845

Storz, G., Wolf, Y. I., & Ramamurthi, K. S. (2014). Small proteins can no longer be ignored. Annu Rev Biochem, 83, 753–777. 10.1146/annurev-biochem-070611-102400

Straub, J., Brenneis, M., Jellen-Ritter, A., Heyer, R., Soppa, J., & Marchfelder, A. (2009). Small RNAs in haloarchaea: Identification, differential expression and biological function. RNA Biol, 6(3), 281–292. http://www.ncbi.nlm.nih.gov/entrez/query.fcgi?cmd=Retrieve&db=PubMed&dopt=Citation&list_uids=19333006

Thauer, R. K. (1998). Biochemistry of methanogenesis: a tribute to Marjory Stephenson. 1998 Marjory Stephenson Prize Lecture. Microbiology (Reading), 144 ( Pt 9), 2377–2406. 10.1099/00221287-144-9-2377

Thauer, R. K., Kaster, A. K., Seedorf, H., Buckel, W., & Hedderich, R. (2008). Methanogenic archaea: ecologically relevant differences in energy conservation. Nat Rev Microbiol, 6(8), 579–591. http://www.ncbi.nlm.nih.gov/entrez/query.fcgi?cmd=Retrieve&db=PubMed&dopt=Citation&list_uids=18587410

Thompson, M. K., Nocedal, I., Culviner, P. H., Zhang, T., Gozzi, K. R., & Laub, M. T. (2022). Escherichia coli SymE is a DNA-binding protein that can condense the nucleoid. Mol Microbiol, 117(4), 851–870. 10.1111/mmi.14877

Thomsen, J., & Schmitz, R. A. (2022). Generating a Small Shuttle Vector for Effective Genetic Engineering of Methanosarcina mazei Allowed First Insights in Plasmid Replication Mechanism in the Methanoarchaeon. Int J Mol Sci, 23(19). 10.3390/ijms231911910

Thomsen, J., Weidenbach, K., Metcalf, W. W., & Schmitz, R. A. (2022). Genetic Methods and Construction of Chromosomal Mutations in Methanogenic Archaea. Methods Mol Biol, 2522, 105–117. 10.1007/978-1-0716-2445-6_6

Tufail, M. A., Jordan, B., Hadjeras, L., Gelhausen, R., Cassidy, L., Habenicht, T., Gutt, M., Hellwig, L., Backofen, R., Tholey, A., Sharma, C. M., & Schmitz, R. A. (2024). Uncovering the small proteome of Methanosarcina mazei using Ribo-seq and peptidomics under different nitrogen conditions. Nat Commun, 15(1), 8659. 10.1038/s41467-024-53008-8

Unoson, C., & Wagner, E. G. (2008). A small SOS-induced toxin is targeted against the inner membrane in *Escherichia coli*. Mol Microbiol, 70(1), 258–270. http://www.ncbi.nlm.nih.gov/entrez/query.fcgi?cmd=Retrieve&db=PubMed&dopt=Citation&list_uids=18761622

Vanderpool, C. K., Balasubramanian, D., & Lloyd, C. R. (2011). Dual-function RNA regulators in bacteria. Biochimie, 93(11), 1943–1949. http://www.ncbi.nlm.nih.gov/entrez/query.fcgi?cmd=Retrieve&db=PubMed&dopt=Citation&list_uids=21816203

Vazquez-Laslop, N., Sharma, C. M., Mankin, A., & Buskirk, A. R. (2022). Identifying Small Open Reading Frames in Prokaryotes with Ribosome Profiling. J Bacteriol, 204(1), e0029421. 10.1128/jb.00294-21

Venkat, K., Hoyos, M., Haycocks, J. R., Cassidy, L., Engelmann, B., Rolle-Kampczyk, U., von Bergen, M., Tholey, A., Grainger, D. C., & Papenfort, K. (2021). A dual-function RNA balances carbon uptake and central metabolism in Vibrio cholerae. Embo j, 40(24), e108542. 10.15252/embj.2021108542

Wagner, E. G. H., & Romby, P. (2015). Small RNAs in bacteria and archaea: who they are, what they do, and how they do it. Adv Genet, 90, 133–208. 10.1016/bs.adgen.2015.05.001

Weidenbach, K., Gutt, M., Cassidy, L., Chibani, C., & Schmitz, R. A. (2022). Small Proteins in Archaea, a Mainly Unexplored World. J Bacteriol, 204(1), e0031321. 10.1128/jb.00313-21

Will, S. (2024). LocARNA 2.0: Versatile Simultaneous Alignment and Folding of RNAs. Methods Mol Biol, 2726, 235–254. 10.1007/978-1-0716-3519-3_10

Will, S., Joshi, T., Hofacker, I. L., Stadler, P. F., & Backofen, R. (2012). LocARNA-P: accurate boundary prediction and improved detection of structural RNAs. Rna, 18(5), 900–914. 10.1261/rna.029041.111

Will, S., Reiche, K., Hofacker, I. L., Stadler, P. F., & Backofen, R. (2007). Inferring noncoding RNA families and classes by means of genome-scale structure-based clustering. PLoS Comput Biol, 3(4), e65. 10.1371/journal.pcbi.0030065

Williams, A. H., Redzej, A., Rolhion, N., Costa, T. R. D., Rifflet, A., Waksman, G., & Cossart, P. (2019). The cryo-electron microscopy supramolecular structure of the bacterial stressosome unveils its mechanism of activation. Nat Commun, 10(1), 3005. 10.1038/s41467-019-10782-0

Yadavalli, S. S., & Yuan, J. (2022). Bacterial Small Membrane Proteins: the Swiss Army Knife of Regulators at the Lipid Bilayer. J Bacteriol, 204(1), e0034421. 10.1128/JB.00344-21

