## Supplementary Files for "Transcription profiling and functional analysis of spRNAs and their corresponding asRNAs in *Methanosarcina mazei*"

### Supplemental data

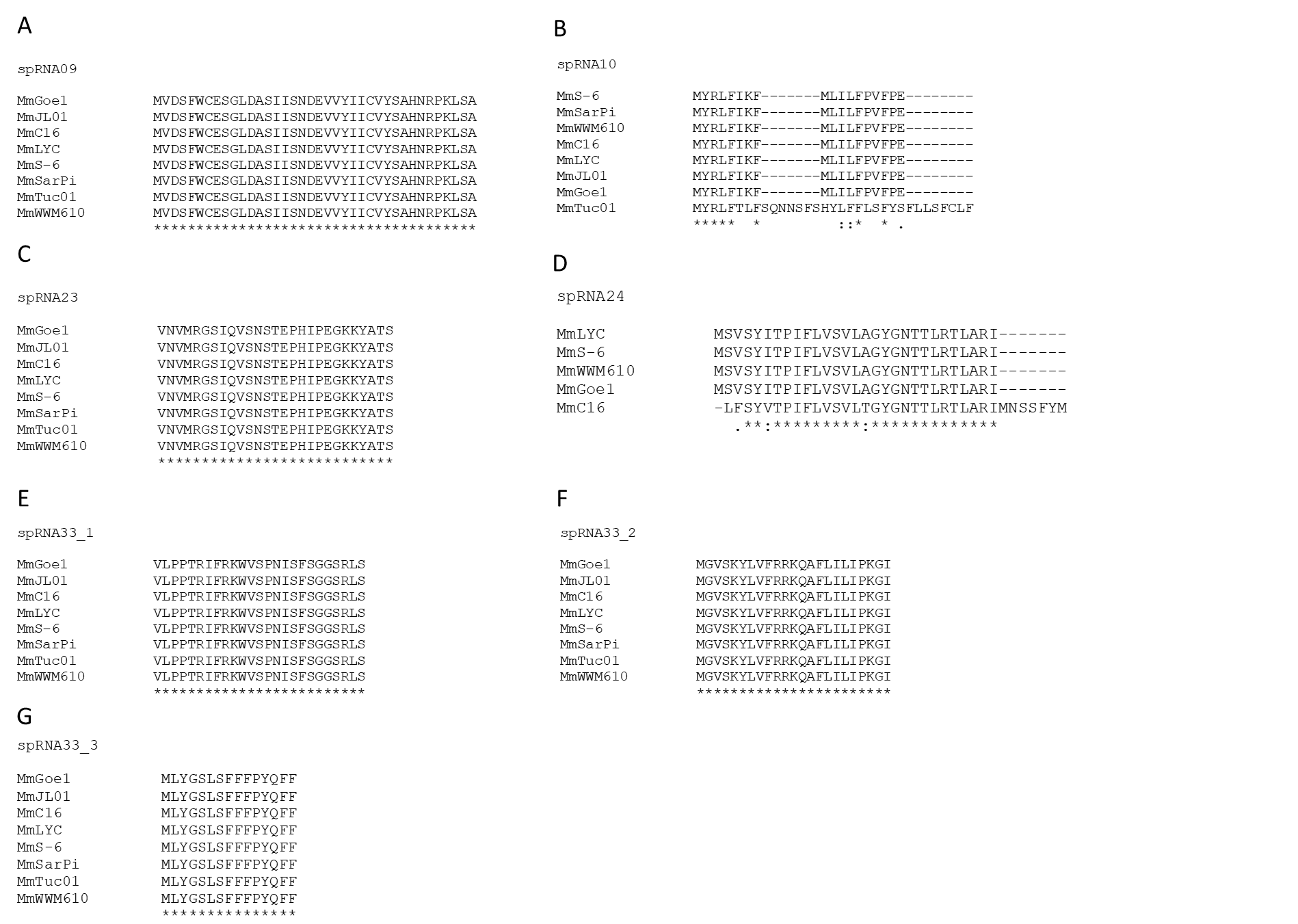

**Figure S1: Amino acid alignment of predicted sORFs**

Mm, *Methanosarcina mazei* strains zm-15, JL01, C16, LYC, S-6, SarPi, WWM610, Tuc01, Goe1

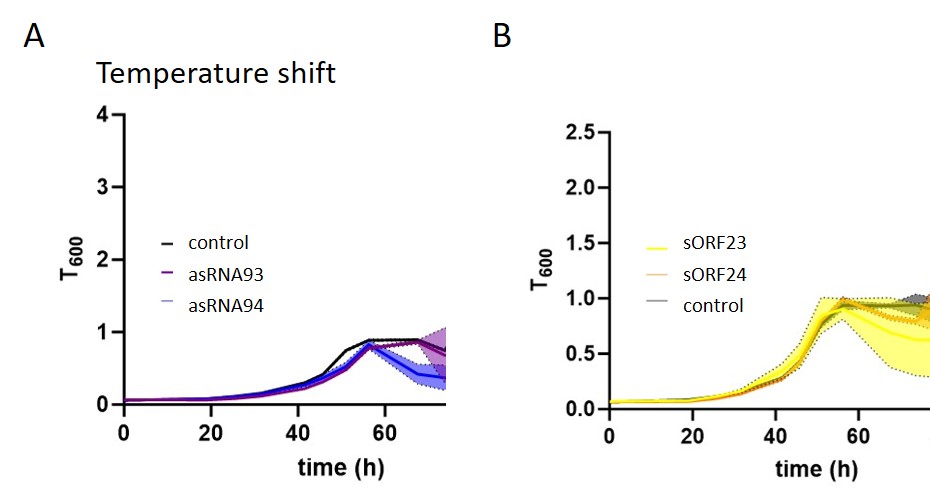

**Figure S2: Growth analysis of *M. mazei* with additional expression of sORF23 or sORF24 and their respective asRNAs under temperature shift.**

*M. mazei* overproduction mutants, expressing (**A**) asRNA93 and asRNA94 or (**B**) sORF23 and sORF24 each under a constitutive promoter as described in Material and Methods, were cultivated in 50 ml at 37°C and shifted to 42° C at T600 ≈ 0.5. As a control *M. mazei* with empty shuttle-vectors pWM321 (control asRNAs) or pRS1595 (control sORFs) were used. The optical density at 600 nm (T_600_) was monitored. The depicted standard deviation (shadows) is based on three individually grown cultures.

**Table S1: Strains used in this study**

| **Organism** | **Description** | **Reference** |
| --- | --- | --- |
| *Methanosarcina mazei* Gö1 | Wild type | DSMZ No. 3647 |
| *M. mazei** | potential cell wall mutant | (Ehlers et al., 2005) |
| *Escherichia coli* DH5α | general cloning strain | (Hanahan, 1983) |
| *E. coli* DH5α λpir | general cloning strain | (Miller and Mekalanos, 1988) |
| *E. coli* K-12 wild type MG1655 | general cloning strain | (Blattner et al., 1997) |

**Table S2: Plasmids used in this study**

| **Plasmid** | **Description** | **Reference** |
| --- | --- | --- |
| pBAD | modified pBAD-TOPO expression plasmid, Amp^R^ | (Unoson and Wagner, 2008) |
| p-sORF24 | pBAD with sORF24, artificial SD sequence and T7 terminator under control of P_BAD_ promoter | This study |
| p-sORF23 | pBAD with sORF23, artificial SD sequence and T7 terminator under control of P_BAD_ promoter | This study |
| pRS1899 | pRS1595 with pmcrB-sORF24, Amp^R^, Pur^R^ | This study |
| pRS1897 | pRS1595 with pmcrB-sORF23, Amp^R^, Pur^R^ | This study |
| pRS1399 | pRS1595 with pmcrB-asRNA24, Amp^R^, Pur^R^ | This study |
| pRS1397 | pRS1595 with pmcrB-asRNA23, Amp^R^, Pur^R^ | This study |
| pWM321 | Shuttle vector for *E. coli* and *M. mazei*, Amp^R^, Pur^R^ | (Metcalf et al., 1997) |
| pRS1595 | Shuttle vector for *E. coli* and *M. mazei*, Amp^R^, Pur^R^ | (Thomsen and Schmitz, 2022) |

**Table S3: Primers used in this study**

bold: restriction site, underlined: artificial 5’ UTR with Shine-Dalgarno sequence, red: mutated sequence

| **Primer** | **Sequence (5’ to 3’)** |
| --- | --- |
| ORF24_1_f | CC**GAATTC**AGAGAAAGAGGAGAAATACTAGatgtcagtttcatatataactcctatttttc |
| ORF24_rev | CA**TCTAGA**CGATTTAGAGCTTGACGGGG |
| F10K_fw | ATAACTCCTATTAAGCTTGTAAGCGT |
| F10K_rev | ACGCTTACAAGCTTAATAGGAGTTAT |
| L11K_fw | AACTCCTATTTTTAAGGTAAGCGTTCTC |
| L11K_rev | GAGAACGCTTACCTTAAAAATAGGAGTT |
| V12K_fw | TCCTATTTTTCTTAAGAGCGTTCTCGC |
| V12K_rev | GCGAGAACGCTCTTAAGAAAAATAGGA |
| G17K_fw | GTTCTCGCAAAATACGGGAATACTACTTTGAG |
| G17K_rev | CCCGTATTTTGCGAGAACGCTTACAA |
| Y18K_fw | CTCGCAGGGAAAGGGAATACTACTTTG |
| Y18K_rev | AGTATTCCCTTTCCCTGCGAGAACGCT |
| T21K_fw | TACGGGAATAAAACTTTGAGAACACTTGC |
| T21K_rev | TCTCAAAGTTTTATTCCCGTACCCTGC |
| T22K_fw | GGGAATACTAAATTGAGAACACTTGCAAG |
| T22K_rev | TGTTCTCAATTTAGTATTCCCGTACCCT |
| sORF24_1 for | ATGTCAGTTTCATATATAACTCC |
| sORF24_1 XhoI rev | **CTCGAG**GGTACCAAAAAATTAAAAG |

**Table S4: Oligonucleotides for northern blot analysis**

| **Name** | **Sequence (5‘- 3‘)** |
| --- | --- |
| spRNA09 | GGTGTGAAAGTGGATTAGATG |
| spRNA10 | ggagttaatgaggcatcttatccac |
| spRNA23 | gacctgtattgaaccccgcataac |
| spRNA24 | gtgttctcaaagtagtattcccgta |
| asRNA28 | GAATGAGGGACGAGTGACCAAC |
| spRNA33 | GATATTTGGAGACACCCATTTGCG |
| asRNA60 | GGTGTGAAAGTGGATTAGATG |
| asRNA61 | gatgcgtaactaaattctacagggc |
| asRNA93 | aaactctaccgaaccccacatac |
| asRNA94 | gggaatactactttgagaacacttgc |
| 5S rRNA | CGCACACTTCAGTACAGTAAGGAA |
